# Glutamate Transporters are Involved in Direct Inhibitory Synaptic Transmission in the Vertebrate Retina

**DOI:** 10.1101/2023.05.01.538911

**Authors:** Stephanie Niklaus, Stella M.K. Glasauer, Peter Kovermann, Kulsum F. Farshori, Lucia Cadetti, Simon Früh, Nicolas N. Rieser, Matthias Gesemann, Christoph Fahlke, Stephan C.F. Neuhauss

**Affiliations:** University of Zurich, Department of Molecular Life Sciences, Winterthurerstrasse 190, CH-8057 Zurich, Switzerland; Institute of Biological Information Processing, Molekular- und Zellphysiologie (IBI-1), Forschungszentrum Jülich, Leo-Brandt-Strasse, DE-52425 Jülich, Germany; EraCal Therapeutics, Wagistrasse 13, 8952 Schlieren, Switzerland; Altos Labs, 1300 Island Drive Redwood City, CA 94065, USA; University of Basel, Department of Biomedizin, Klingelbergstrasse 50 4056 Basel, Switzerland

## Abstract

In the central nervous system of vertebrates, glutamate serves as the primary excitatory neurotransmitter. However, in the retina, glutamate released from photoreceptors causes hyperpolarization in postsynaptic ON-bipolar cells through a glutamate-gated chloride current, which seems paradoxical. Our research reveals that this current is modulated by two excitatory glutamate transporters, EAAT5b and EAAT7. These transporters are located at the dendritic tips of ON-bipolar cells and interact with all four types of cone photoreceptors. The absence of these transporters leads to a decrease in ON-bipolar cell responses, with eaat5b mutants being more severely affected than eaat5b/eaat7 double mutants, which also exhibit altered response kinetics. Biophysical investigations establish that EAAT7 is an active glutamate transporter with a predominant anion conductance. Our study is the first to demonstrate the direct involvement of postsynaptic glutamate transporters in inhibitory direct synaptic transmission at a central nervous system synapse.

## Introduction

The first visual synapse encompasses connections between presynaptic photoreceptors and postsynaptic bipolar and horizontal cells. Bipolar cells vertically relay the signal they receive from photoreceptors to ganglion cells, which constitute the retinal output neurons. In the photoreceptor synapse, the light induced signal is transmitted from photoreceptors into two parallel pathways, the ON and OFF pathway mediated by two types of bipolar cells, the ON- and OFF-bipolar cells, respectively. While ON-bipolar cells depolarize upon light increments, OFF-bipolar cells depolarize in darkness. This separation is based on the differential expression of glutamate receptors. OFF bipolar cells are depolarized by glutamate via AMPA/kainite receptors. Photoreceptors tonically release glutamate in darkness, which binds to the class III metabotropic glutamate receptor mGluR6 on ON-bipolar cell dendrites. This in turn causes the activation of a G-protein signaling cascade leading to the closure of the constitutively open cation conducting ion channel TRPM1 (Morgans et al. 2009; Morgans et al. 2010). In incremental light, glutamate release by photoreceptors is reduced, rendering the mGluR6 signaling cascade inactive. The consequential opening of TRPM1 causes ON-bipolar cells to depolarize. This mGluR6-TRPM1 signaling cascade is evolutionarily conserved in vertebrates (e.g. (Morgans et al. 2009; Huang et al. 2012)). However, the presence of a second glutamatergic input mechanism, which is additionally involved in ON-bipolar cell activation has been first suggested in teleost fishes and may well be present in vertebrates up to mammals. First experimental evidence of this second glutamatergic input mechanism dates back to 1979 when Saito *et al*. showed a resistance increase in carp ON-bipolar cells at photopic conditions and a decrease thereof at scotopic conditions (Saito et al. 1979). A series of studies on different teleost retinas later demonstrated the presence of a glutamate-activated chloride current in ON-bipolar cells (Grant and Dowling 1995, 1996; Connaughton and Nelson 2000; Wong et al. 2005a; Wong et al. 2005b; Thoreson and Miller 1993). The sensitivity of this current to the non-specific glutamate transporter blocker DL-threo-beta-Benzyloxyaspartate (TBOA) suggested the current to be mediated by excitatory amino acid transporters (EAATs) (Thoreson and Miller 1993; Wong et al. 2005a; Wong et al. 2005b). EAATs are glutamate transporters and are members (together with the neutral amino acid transporters) of the solute carrier 1 (SLC1) family (Arriza et al. 1993; Shafqat et al. 1993) (Kovermann et al. 2021). These transporters are pre- and post-synaptically expressed on neurons and glia cells. The transport of glutamate from the synaptic cleft into the cell by EAATs is electrogenic. Transmembrane concentration gradient of the co-transported Na^+^ is the transporters driving force with one molecule of glutamate being co-transported with 3 Na^+^ and one H^+^ followed by a counter-transport of one K^+^ (reviewed by (Fahlke et al. 2016)). Interestingly, substrate binding generates a thermodynamically uncoupled, Cl^-^ conductance through EAATs with current amplitudes highly varying between different EAAT isoforms (Fairman et al. 1995a; Mim et al. 2005; Palmer et al. 2003) (Schneider et al. 2014; Torres-Salazar and Fahlke 2007). In essence, EAATs are bifunctional, serving as transporter, as well as anion channels and thus are capable of modulating signal transmission by uptake of glutamate from the synaptic cleft and modulating the cell membrane potential. EAAT5 has been proposed to mediate the ON-response due to its biophysical characteristics (Thoreson and Miller 1993). Both, human and mouse EAAT5 have been extensively characterized in isolated expression systems. EAAT5 is a low capacity transporter with a transporter current (generated by the co- and counter-transported ions) below resolution threshold (Gameiro et al. 2011; Schneider et al. 2014). However, EAAT5 was shown to be associated with a large voltage-dependent anion conductance comparable to a bona fide Cl^-^ channel, which optimally functions at negative potentials (Gameiro et al. 2011; Schneider et al. 2014) (Gehlen et al. 2021, 2021).

Teleosts still have the complete ancestral vertebrate complement of 7 EAAT genes. Moreover EAAT1, 2, 5 and 6 even have two ohnologs stemming from an ancient whole genome duplication (Gesemann et al. 2010).

Here we analyzed the expression patterns of the two ON-bipolar cell glutamate transporters EAAT5b and EAAT7. We show that both transporters are co-expressed on ON-bipolar cell dendritic tips contacting all cones. Functional analysis on knockout (KO) animals showed a reduced b-wave response in the electroretinogram (ERG) of *eaat5b* mutants, which is larger in *eaat5b/eaat7* double mutants. Intriguingly, the implicit time (time to peak of the b-wave) is shorter in *eaat5b* single and *eaat5b/eaat7* mutants. Hence, our study provides evidence of EAAT5b and EAAT7 generating hyperpolarizing currents in darkness with different kinetics. This is the first report directly showing that glutamate mediated EAAT hyperpolarizing currents are involved in direct synaptic transmission in a central nervous synapse.

## Material and Methods

### Zebrafish husbandry

Zebrafish (*Danio rerio*) of the wildtype (WT) strains Wik and Tübingen were kept and bred under standard condition at 28°C in a 14/10 h light/dark cycle as described before (Mullins et al. 1994). Embryos and larvae were raised until 5 days post fertilization (dpf) in E3 medium containing methylene blue or in 1-phenyl-2-thiourea (PTU, Sigma-Aldrich) if prevention of pigment formation was required. From 5 dpf on, larvae were raised in facility water.

The following transgenic lines were used for immunohistochemistry: *Tg(zfSWS1–5.5A:EGFP)* (Takechi et al. 2003),, *Tg(zfSWS2–3.5A:EGFP)* (Takechi et al. 2008), *Tg(zfRh2-2:EGFP)* (Tsujimura et al. 2007), *Tg(zfLWS:EGFP)* (Tsujimura et al. 2010) and *Tg(zfRh1-3:EGFP)* (Hamaoka et al. 2002), expressing EGFP in UV, blue, green, red cones and rods, respectively.

### Generation of CRISPR/Cas9 knockout lines

CRISPR targets were selected using the target site prediction tools www.zifit.partners.org and https://chopchop.rc.fas.harvard.edu. The following genomic target sites (GGN18) were chosen: 5’-GGTGGTGGTGGGAATCGTCA-3’ on exon 3 of *eaat5b* and 5’-GGGAACCCAAAACTCAGGTC-3’ on exon 3 of *eaat7*. sgRNAs were prepared using a PCR based protocol as described by (Gagnon et al. 2014) with minor adaptations. Briefly, double strand DNA template oligos were synthesized using a Phusion High-Fidelity DNA Polymerase (New England BioLabs) with the T7 specific forward primer 5’-GAAATTAATACGACTCACTATA**GGN18**GTTTTAGAGCTAGAAATAGC-3’ together with the common reverse primer 5’- AAAAGCACCGACTCGGTGCCACTTTTTCAAGTTGATAACGGACTAGCCTTATTTTAACTT GCTATTTCTAGCTCTAAAAC-3’. The generated amplicon was purified after gel electrophoresis using the QIAquick Gel Extraction Kit (Qiagen) and served as template for *in vitro* transcription with MEGAshortscript T7 Transcription Kit (Ambion). Purification of synthesized sgRNA was carried out using the Megaclear Kit (Ambion) which was followed by an ethanol precipitation.

The CRISPR injection mix containing 814 ng/µl GFP-tagged Cas9 protein (kindly provided by Prof. Dr. C. Mosimann and Prof. Dr. M. Jinek), 150 ng/µl sgRNA and 300 mM KCl (Burger et al. 2016) was incubated at 37°C for 10 min prior injection allowing protein-sgRNA complex formation. 1 nl of the mix was injected into the cell of a one-cell stage embryo using a FemtoJet Microinjector (Eppendorf).

Mosaic F0 founder fish were outcrossed to WT fish and resulting heterozygous F1 generation was genotyped (PCR amplification of lysed fin biopsies, cloning and sequencing of plasmids) using the following primers: 5’-TTGATGTCAGGTTTGGCG-3’ and 5’-TGATGGGTTTTCCGCTGT-3’ for *eaat5b* and 5’-ATGTCCACCACAGTAATCG-3’ and 5’-TTGAAAACAGGCCTGGAC-3’ for *eaat7*. Fish carrying a +79 bp indel in *eaat5b* and a −4 bp deletion in *eaat7* were used for all experiments.

*eaat5b* and *eaat7* mutant fish were genotyped by PCR amplification of the genomic region using the primers listed above. Amplicon length revealed the genotype of *eaat5b*, with a WT amplicon of 180 base pairs (bp) and a mutant fragment of 259 bp. *eaat7* PCR amplicon was digested using DdeI resulting in two WT fragments of 116 bp and 73 bp and a sole mutant fragment of 185 bp. *eaat5b* and *eaat7* double KO fish were generated by crossing *eaat5b*^-/-^ to *eaat7*^-/-^ fish.

### Cloning of *eaat* genes and mRNA *in situ* hybridization

Fragments of *eaat* genes were PCR amplified using a JumpStart Taq polymerase (Sigma-Aldrich) and the primer pairs *eaat5b*_135_s 5’-GGAGCAGGAAGTCAAGTA-3’ with *eaat5b*_600_as 5’-GTCCGTCCCATTATCGTC-3’ and *eaat7*_142_s 5’-GTAATAGCAGGCACAGTGATG-3’ with *eaat7*_1001_as 5’-CCCAGAGCAGTGATCCAAG-3’. *In situ* probes were prepared according to the protocol described by (Niklaus et al. 2017). *In situ* hybridization was performed on 5 dpf PTU treated whole mount larvae and sections of adult retinae. Adult eyes and whole larvae were fixed with 4% paraformaldehyde (PFA) overnight at 4°C. Eyes were cryoprotected in 30% sucrose (in phosphate buffered saline (PBS)) overnight at 4°C, embedded in Tissue-Tek O.C.T. Compound (Sakura Finetek) and sectioned at 16 μm. Both, whole mount and slide *in situ* hybridization were carried out based on the protocol published by (Thisse and Thisse 2008) with adaptations described in (Huang et al. 2012). Hybridization and stringency washes were carried out at 64°C for both genes.

### Generation of Antibodies

Zebrafish peptide specific antibodies were generated in an 87-day classical program by Eurogentec S.A. (Seraing, Belgium). Guinea pigs and rabbits were immunized with the EAAT5b epitope N-PDRKKPPVPPRHLKHRDKDHCA-C and EAAT7 epitope N-KLRSGQVSSAPRNQEV-C, respectively. Guinea pig anti-EAAT5b and rabbit anti-EAAT7 antibodies were column purified by Eurogentec.

### Immunohistochemistry

Eyes of adult fish were fixed with 4% PFA for 30 min at room temperature (RT) or with 2% trichloroacetic acid (TCA) for 25 min at RT. Fixed eyes were washed twice with PBS prior to dehydration in 30% sucrose (in PBS) overnight at 4°C. Eyes were embedded in Richard-Allan Scientific Neg-5 Frozen Section Medium (Thermo Fisher Scientific) and sectioned at 16 μm. Sections mounted on superfrost coated slides (Thermo Fisher Scientific) were dried for 30 min a 37°C before the staining. Sections were washed with PBS and blocked for 45 min with blocking solution (10% normal goat serum, 1% bovine serum albumin, 0.3% Triton in PBS, pH 7.4). Primary antibodies diluted in blocking solution were applied ON at 4°C. The following antibodies were used: guinea pig anti-EAAT5b (1:100), rabbit anti-EAAT7 (1:400), rabbit anti-mGluR6b (1:750) (Huang et al. 2012), mouse anti-PKCα MC5 (Novus Biologicals, NB200-586) (1:500), mouse anti-Synaptic Vesicle 2 (SV2) (DSHB, Iowa, USA) (1:100), chicken anti-GFP (A20162; Invitrogen) (1:500) and rabbit anti-PKCβC16 (#E1313, Santa Cruz Biotechnology) (1:500). Primary antibodies were detected with the following secondary antibodies (all 1:500 in PBS) for 1.5 h at RT: goat anti-rabbit, goat anti-guinea pig, goat anti-chicken and goat anti-mouse all conjugated to Alexa Fluor (AF) 488, 568 or 647 (Invitrogen, Molecular Probes). For stimulated emission depletion (STED) microscopy, goat anti-guinea pig AF 488 and goat anti-rabbit Atto 594 secondary antibodies were used. To counterstain green fluorescence, Bodipy TR methyl ester (Thermo Fisher Scientific) diluted 1:300 in PDT (0.1% Triton-X100 and 1% DMSO in PBS) was applied for 20 min to sections after washing secondary antibodies. Confocal laser scanning imaging was done with a TCS LSI confocal microscope (Leica) and STED images were taken with a CLSM-Leica SP8 inverse STED 3X microscope. The same confocal settings were used for imaging corresponding stainings on mutant and WT animals.

### Retinal histology

Prior embedding in Technovit 7100 (Kulzer Histotechnik) 5 dpf larvae and adult eyes were fixed in 4% PFA overnight at 4°C. Embedding, sectioning and staining of histological sections was done according to the protocol described by (Niklaus et al. 2017). Images were taken in the bright field mode of a BX61 microscope (Olympus).

#### White light and flicker electroretinography

White light electroretinograms (ERG) were recorded on 5 dpf WT and mutant animals in a double blinded manner. Larvae were dark adapted for at least 30 min and preparations prior recordings were done under a dim red light to prevent bleaching of photo pigment. The larval eye was removed and placed on a filter paper on top of an agarose gel. The reference electrode was inserted into the agarose gel and the recording electrode, a glass capillary GC100-10 (Harvard Apparatus, Holliston USA) with a tip diameter of 20-30 μm filled with E3 was placed on top of the cornea.

##### White light ERG

A series of 5 white light stimuli of increasing light intensities (log −4 to log 0) were presented to the eyes. Duration of stimuli was 100 ms with inter stimulus intervals of 7 sec. For *eaat5b* mutant recordings, 100% light (log 0) corresponds to 6800 lux or 75 W/m^2^ (Zeiss XBO 75 W light source). For recordings of *eaat7*-/- and double mutants, log 0 corresponds to 696 lux (high power xenon light source HPX-2000, Ocean Optics) if measured with a TES 1335 Light Meter.

##### Flicker ERG

The flicker fusion frequency (FFF), defined as the frequency of visual stimuli that cannot be temporally resolved anymore was assessed as a read-out for temporal resolution of vision. 15 ms light stimuli were presented to the eyes at different frequencies starting at 7 Hz and increasing in 1 Hz steps. ERG responses to each frequency were recorded for 2 sec.

### Analysis of ERG data

ERG traces were analyzed using Excel and Igor-Pro Software (Wave Metrics). B-wave amplitudes were analyzed as a proxy for ON-bipolar cell depolarization. The first 50 ms of each recording were averaged and taken as baseline values. B-wave amplitudes were statistically compared between WT and mutant animals by a two tailed t-test using IBM SPSS Statistics 22.

The b-wave implicit time was analyzed as a read-out for transporter kinetics. The implicit time is defined as the time from light onset to the onset of the b-wave. By extrapolating a line that goes through two points on the curve at Y=80% and Y=20% of the b-wave amplitude, respectively, the implicit time was defined as the intersection of this line and the X-axis. Values below 50 ms and above 230 ms were filtered out. Implicit times were statistically compared between WT and mutants with a two tailed t-test using IBM SPSS Statistics 22.

Frequency components of the FFF ERG were assessed by a Fast Fourier Transform using MATLAB R2016a. Statistical comparison of FFFs of mutant and WT animals was carried out with a two tailed t-test using IBM SPSS Statistics 22.

All ERG results are plotted in box-and-whisker plots with boxes reaching from the first to the third quartile, the line within the box representing the median and top and bottom whiskers reaching to the maximum or minimum of obtained values, respectively.

### Image processing and assembly

Images were processed with Imaris (Bitplane) and Adobe Photoshop CC 2017 and assembled using Adobe Illustrator CC 2017.

### Quantitative reverse transcription PCR

5 dpf larvae were anesthetized on ice and eyes dissected using an insect pin and a syringe needle in a dish containing RNA later (Sigma-Aldrich). To confirm genotype, remaining tissue was lysed and gDNA amplified by means of PCR (KAPA2G Fast HotStart PCR kit, KAPA Biosystems) as described above. Total RNA of pools of larval eyes (18-28 per sample) was extracted using the RNeasy Plus Mini kit (Qiagen). RNA was reverse transcribed to cDNA using the Super Script III First-strand synthesis system (Invitrogen) using 1:1 ratio of random hexamers and oligo (dt) primers.

qPCR reactions were performed using SsoAdvanced Universal SYBR Green Supermix on a CFX96 Touch Real-Time PCR Detection System (Bio-rad). Primer efficiencies were calculated by carrying out a dilution series. After the primer efficiencies were determined equal, eye samples were used for qPCR using 1 ng of cDNA per reaction. The controls “no reverse transcription control” (nRT control) and “no template control” (NTC) were performed with every qPCR reaction. *rpl13a* was chosen as a reference gene. All reactions were performed in technical triplicates. Data was analyzed in CFX Maestro Software from Bio-Rad, Microsoft Excel and R. Statistical analysis was performed on the log transformed normalized expression per sample.

Primers used were *eaat7* (GCCCACTCACGACAACCAG; ATCTCGTTCCCGTTCCCTCAG; amplicon size 133bp), *eaat5b* (GCTGATGCGCATGTTGAAGATG; GACGACTGGAACACTTGGCATC; 95bp), and *rpl13a* (TCTGGAGGACTGTAAGAGGTATGC; AGACGCACAATCTTGAGAGCAG; 148bp).

### Heterologous expression of EAAT-fusion proteins in mammalian cells

Transient transfection of HEK293T (Invitrogen) with the EAAT-YFP fusion proteins *dr*EAAT7, *r*EAAT4, and *h*EAAT1 in pcDNA3.1 vectors (Invitrogen) was performed using the Ca^2+^phosphate technique as previously described (Chahine et al. 1994).

### Confocal microscopy of EAAT7 fusion proteins

Confocal imaging was carried out on living cells grown on poly-lysine coated *µ*-dishes (ibidi) without fixation using an inverted confocal laser scanning microscope (Leica TCS SP5, Leica Microsystems, Heidelberg, Germany) equipped with a 63×/1.4 oil immersion objective. YFP was excited using a 488 nm laser and emission light was recorded between 525 and 540 *nm*. Spatial distributions of YFP-fluorescences were analyzed with Fiji package (NIH).

### Electrophysiology

Standard whole-cell patch clamp recordings were performed using a Axoclamp 200B (Molecular Devices) amplifier. Borosilicate glass electrodes were pulled and used with resistances between 1 - 2.5 MΩ. Compensation of 80 % of the series resistance reduced voltage-errors. Recorded currents were filtered at 1 kHz and digitized by a Digidata 1550A (Molecular Devices) digitizer at a sampling rate of 10 kHz. To record anion currents, currents were measured using standard external solution containing in the bath (in mM): 140 NaNO_3_, 4 KCl, 2 CaCl_2_, 1 MgCl_2_, 0.1 L-glutamate, and 10 HEPES/NaOH, pH 7.4, and in the pipette: 110 KNO_3_ or NaNO_3_, 2 MgCl_2_, 5 EGTA, 10 HEPES (KOH or NaOH), pH 7.4. For measurements of glutamate transport, all anions were equimolar replaced by the impermeable corresponding gluconate salt. All experiments were carried out using agar bridges with 2 M KCl in 1 % Agarose to connect the Ag/AgCl electrode. Liquid junction potential differences were calculated in pClamp 10.5 (Molecular Devices) and protocols adjusted a priori. Currents were analyzed using Clampfit (Molecular Devices) and Sigma Plot (SysStat) software. Current-voltage relationships show point plots of mean current amplitudes with standard errors (SE). Box plots of current amplitudes at a single holding potential span the first to the third quartiles, with whiskers reaching maximum and minimum values, respectively. The lines indicate the median values. Pair-wise comparison of data sets was done with *one-way* ANOVA and Holm-Sidak *post hoc* testing.

## Results

### EAAT5b and EAAT7 are expressed on ON-bipolar cell dendritic tips

In order to identify EAATs that are potentially involved in synaptic transmission at the photoreceptor synapse, we cloned all 11 orthologues in zebrafish and performed an mRNA in situ expression survey for expression in second order neurons in the retina (Ref Gesemann, unpublished data). For two orthologues, *eaat5b* and *eaat7*, we found expression in the inner nuclear layer of the retina making them candidates as mediators in direct synaptic transmission.

In the larval retina, the transcript of *eaat5b* is exclusively found in the inner nuclear layer (INL) (Figure 1A). During adult stages, *eaat5b* mRNA is additionally expressed in the outer nuclear layer (ONL) (Figure 1B). *eaat7* message is found in cells of the INL, both in the 5 dpf larvae and in the adult retina (Figure 1C, D). Furthermore, both *eaat5b* and *eaat7* mRNA are additionally detected in other regions of the central nervous system, which however is not subject of this study.

**Figure 1:**
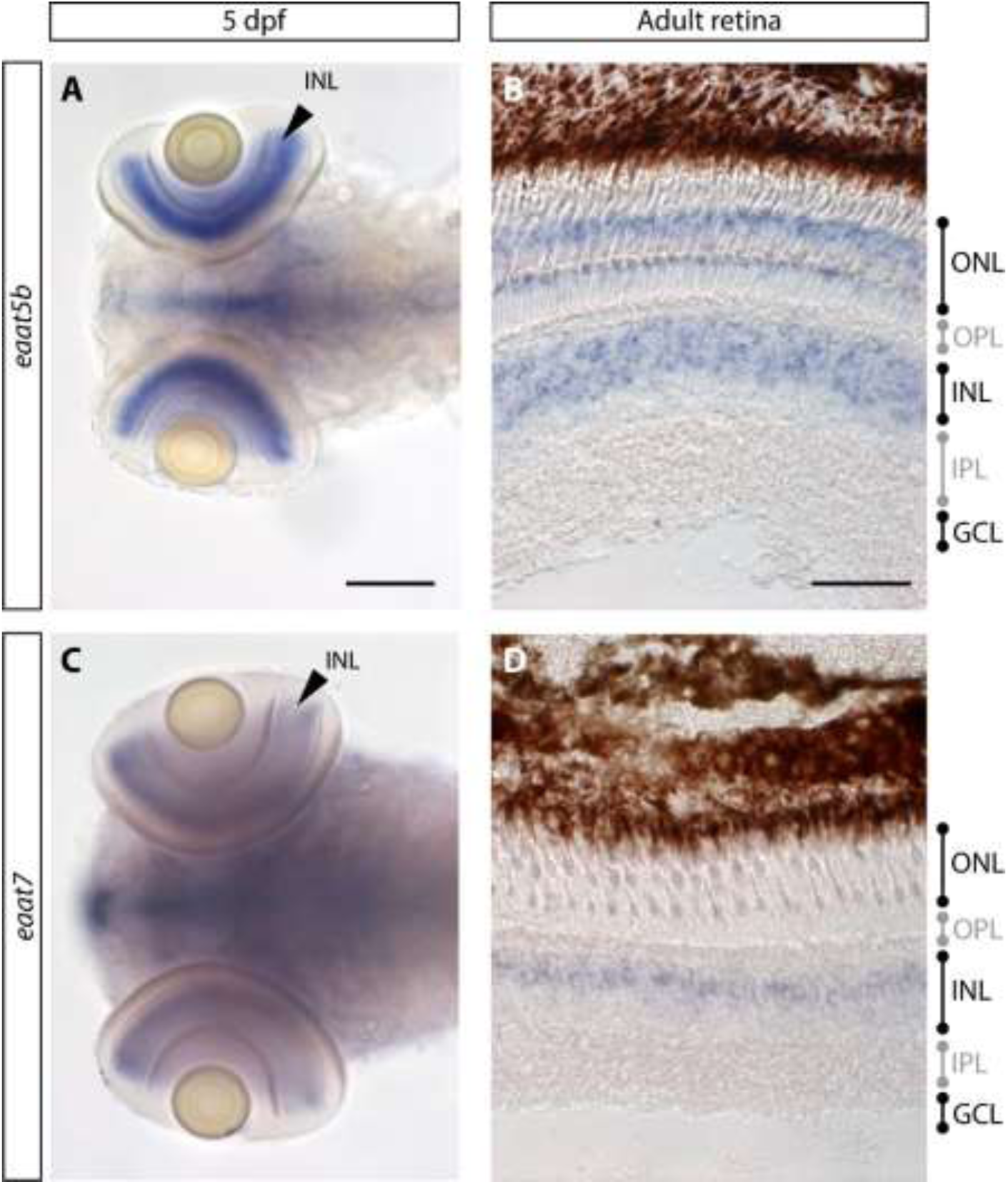
*eaat5b* and *eaat7* mRNA is detected in the inner nuclear layer (INL) of the retina. (A, B) The transcript of *eaat5b* is found in the INL both at larval and adult stages. (B) In the adult retina, *eaat5b* mRNA is further detected in the outer nuclear layer. (C) *eaat7* transcript in the retina is exclusively found in the INL in larvae as well as in the adult eye (D). Scale bar is 100 μm in (A,C) and 50 μm in (B, D). PhR: photoreceptor; OPL: outer plexiform layer; INL: inner nuclear layer, IPL: inner plexiform layer; GCL: ganglion cell layer.

In order to assign the expression to specific inner retinal cell types and to access subcellular localization, we generated paralog specific polyclonal antibodies. These antibodies were used for immunohistochemical stainings. Both transporters share a common dotted expression pattern in the outer plexiform layer (OPL). Co-staining of EAAT5b and EAAT7 with the ON-bipolar cell marker PKCα reveals that both transporters localize to dendritic tips of ON-bipolar cells (Figure 2C, I). EAAT5b and EAAT7 are always co-expressed within a dendritic tip (Figure 2A, B). Surprisingly, super-resolution microscopy reveals adjacent but not overlapping localization within a dendritic terminal (Figure 2B). While EAAT5b localizes more proximal to the synaptic cleft, EAAT7 is located slightly more distal to it. Next, we asked which photoreceptor subtypes are contacting the EAAT5b/7 positive synapses. Therefore we used a number of transgenic lines expressing GFP driven by different opsin promoters, namely *Tg(zfSWS1–5.5A:EGFP)* (Takechi et al. 2003), *Tg(zfSWS2–3.5A:EGFP)* (Takechi et al. 2008), *Tg(zfRh2-2:EGFP)* (Tsujimura et al. 2007), *Tg(zfLWS:EGFP)* (Tsujimura et al. 2010) and *Tg(zfRh1-3:EGFP)* (Hamaoka et al. 2002). Immunolabeling on retinal sections of these transgenic lines show that EAAT5b as well as EAAT7 proteins localize to ON-bipolar cell dendrites contacting all four cone subtypes (UV-, blue-, green- and red light sensitive cones) but only a minority of rod spherules (Figure 2 D-H’ and J-N’). Taken together, these data implies that these two transporters participate in cone signal transmission.

**Figure 2:**
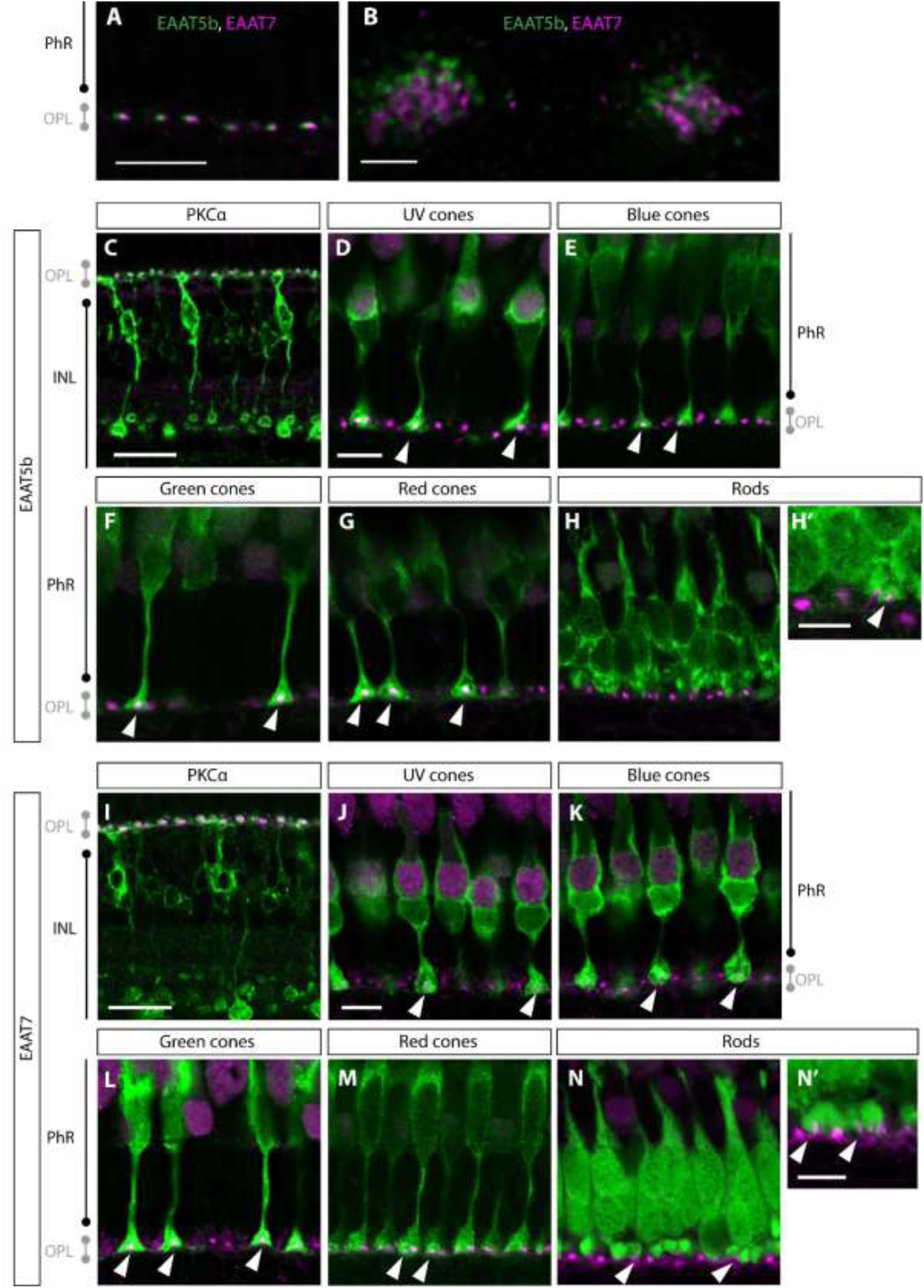
EAAT5b and EAAT7 co-localize on ON-bipolar cell dendritic tips. (A, B) Adult retinal sections stained with anti-EAAT5b (green) and anti-EAAT7 antibody (magenta) shows punctate co-localization of EAAT5b and EAAT7 in the outer plexiform layer. (C) Immunostaining of EAAT5b (magenta) and PKCα (green) demonstrate the dendritic expression of EAAT5b on ON-bipolar cells. EAAT5b localizes to ON-bipolar cells contacting UV (D), blue (E), green (F) and red (G) cones but only a minority of rods (H and H’). Similarly, EAAT7 (magenta) is specific to ON-bipolar cell (stained with PKCα in green) dendrites (I) and localizes to UV (J), blue (K), green (K) and red (M) cone- and few rod synapses (N and N’). Scale bar in A is 15 µm, scale bar in B is 1 µm, scale bar in C is 20 µm, scale bar in D is 8 µm om D, E-H, scale bar in H’ is 5 µm, scale bar in I is 20 µm, scale bar in J K-N is 8 µm, scale bar in N’ is 5 µm. PhR: photoreceptor; OPL: outer plexiform layer; INL: inner nuclear layer.

### Loss of EAAT5b and EAAT7 does not affect retinal morphology and synaptic structure

In an effort to elucidate the function of these transports/receptors, we generated knock-out mutant strains using CRISPR/Cas9 genome editing technology. Both generated genome modifications are predicted to result in severely truncated proteins of no function (Supplemental Figure 1). As both antibody binding epitopes are either C-terminal of the predicted stop codon (EAAT5b) or disrupted by the deletion (EAAT7) (Supplemental Figure 1), neither EAAT5b nor EAAT7 was detected in the corresponding mutants. This provides additional evidence for the paralog specificity of the generated antibodies.

Homozygous mutation of *eaat5b* does not result in any obvious alterations in staining abundance and localization of EAAT7 (Figure 3B) and vice versa, *eaat7* KO animals show an undisrupted distribution of EAAT5b (Figure 3D). In order to provide a more quantifiable measure, we performed quantitative PCR on both mutant strains and indeed found no changes in *eaat5b* mRNA abundance in *eaat7*-/- mutants and vice versa of *eaat7* mRNA in *eaat5b*-/- mutant larvae (Supplemental Figure 2). Combined loss of both transporters, EAAT5b and EAAT7, has no influence on synaptic structure nor localization of synaptic proteins such as the pre-synaptic marker Synaptic Vesicle 2 (SV2) or the post-synaptically expressed metabotropic glutamate receptor 6b (mGluR6b) (Figure 3 I-J). Additionally, morphology of ON-bipolar cells visualized by PKCα immunofluorescence appears normal in double mutants (Figure 3 K-L).

**Figure 3:**
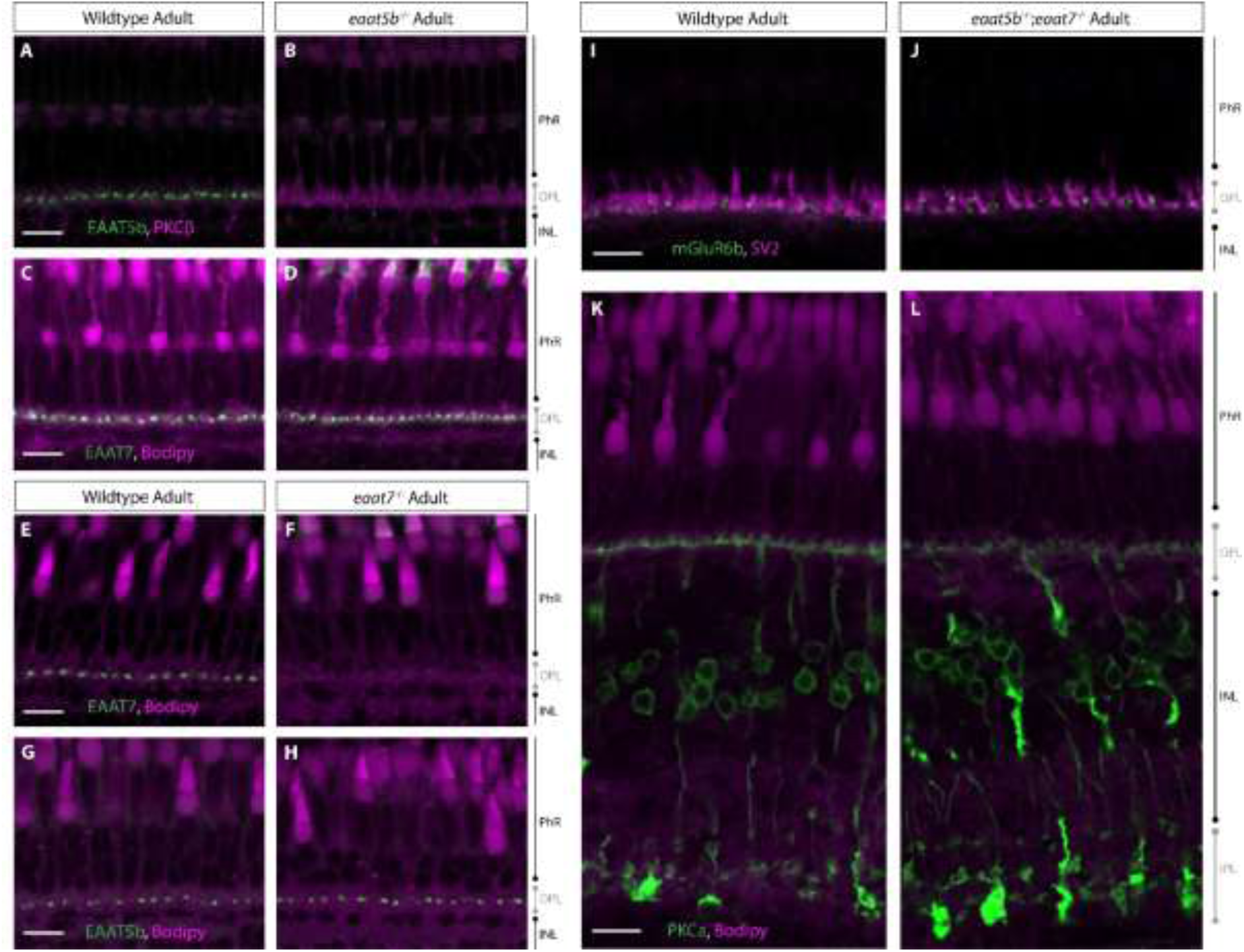
Normal retinal morphology in *eaat5b*-/-, *eaat7*-/- and double KO animals. Immunofluorescent staining of EAAT5b (green) on WT (A) and *eaat5b* mutant retina (B) reveals the absence of EAAT5b protein in mutants. Mutation in *eaat5b* does not affect localization and abundance of EAAT7 (green), seen by comparable immunofluorescence in WT (C) and *eaat5b* mutants (D). EAAT7 antibody staining on WT (E) and *eaat7* KO retina (F) reveals no EAAT7 protein (green) immunofluorescence can be detected in *eaat7*-/- retina (F), however, expression pattern of EAAT5b in *eaat7* KO animals (H) is not affected (if compared to WT (G)). Synaptic structure as well as ON-bipolar cell morphology and abundance in double KO animals are not affected. Both, the localization of pre-synaptic marker SV2 (magenta) as well as the post-synaptic mGluR6b (green) are not altered in the retina of *eaat5b*-/-;*eaat7*-/- (J) if compared to WT (I). Furthermore, shape of ON-bipolar cells stained with PKCα (green) is comparable between double mutants (L) and WT (K). Scale bars in A, C, E, G, I and K are all 10 μm and also correspond to B, D, F, H, J and L, respectively.

Retinal morphology of 5 dpf larvae and adult eyes of homozygous mutant was assessed on histological plastic sections. Loss of either EAAT5b or EAAT7 does not cause any retinal morphology changes. Lamination as well as thickness of the retinal layers appears normal in the KO animals (Supplemental Figure 3 C-F). Furthermore, also double KO animals (*eaat5b*-/-;*eaat7*-/-) did not reveal any morphological alterations of the retina, neither at larval nor at adult stages (Supplemental Figure 3 G-H).

In summary, neither retinal morphology alterations nor changes in synaptic protein distribution and abundance are apparent in single and double mutants, indicating that any functional phenotype does not arise from structural changes but would be solely attributable to functional alterations of the corresponding transporter.

### EAAT5b and EAAT7 contribute to the ON-response

Next, we assessed outer retinal function in the mutants. We employed electroretinography (ERG), providing a ready assessment of outer retinal function. We quantified the b-wave in larval retinae as a read-out of ON-bipolar cell depolarization. While the loss of EAAT5b causes a subtle decrease in the b-wave amplitude at medium light intensities (log −3 and log −2), the b-wave amplitudes for dim light (log −4) and bright light (log −1 and log 0) do not significantly differ between WT and *eaat5b* KO animals (Figure 4A). Homozygous mutant larvae for *eaat7* show no phenotype in the ERG upon white light stimulation at all light intensities tested (log −4 to log 0) (Figure 4B).

**Figure 4:**
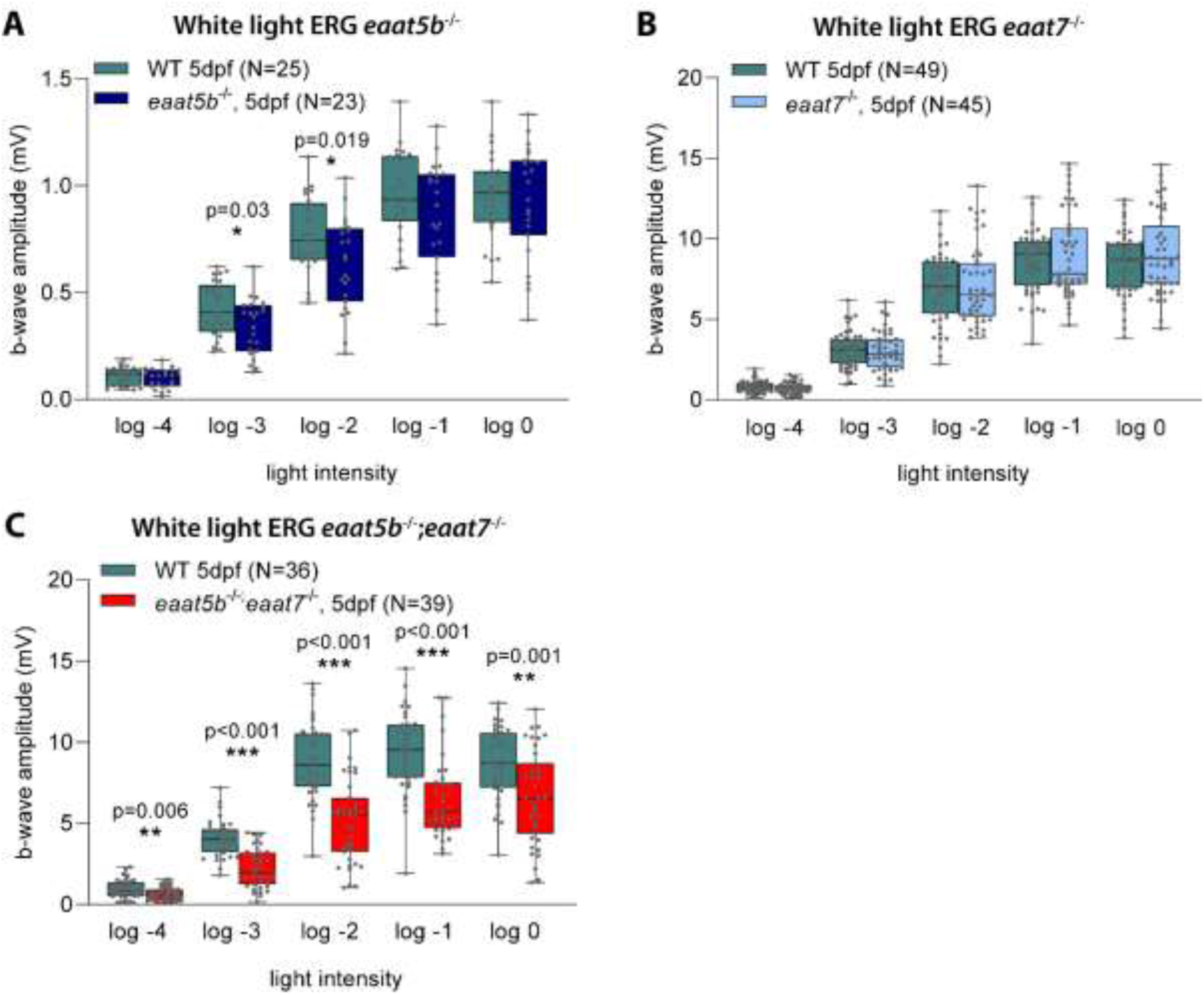
Involvement of EAAT5b and EAAT7 in ON-bipolar cell activation. ERG was recorded on *eaat5b*-/- (A), *eaat7*-/- (B) and *eaat5b*-/-;*eaat7*-/- (C) animals. B-wave amplitudes are plotted in box-and-whisker plots. (A) Loss of EAAT5b causes a small, slightly significant decrease in the b-wave amplitude for medium light intensities (log −3 and log −2). (B) Homozygous mutation in *eaat7* does not affect ON-bipolar cell activation in a by white light ERG measurable way. (C) Double KO animals show a significant reduction in the b-wave amplitudes across the 5 irradiances. Note that the data plotted in panel A was recorded at lower light intensities than in panels B and C.

However, double KO larvae lacking both EAAT5b and EAAT7 reveal a strong ERG phenotype. B-wave amplitudes of *eaat5b*-/-;*eaat7*-/- are strongly reduced across all light intensities tested, indicating a redundant involvement of these two transporters in ON-bipolar cell activation (Figure 4C).

We next investigated the kinetics of signal transmission in these mutants. Interestingly, loss of the two ON-bipolar cell glutamate transporters differentially affects kinetics of signal transmission in the first visual synapse. The b-wave implicit time, defined as the time from the onset of the light stimulus to the onset of the b-wave is significantly decreased in *eaat5b* mutant animals compared to WT (Figure 5A). This demonstrates that the ON-response commences faster in animals without a functional EAAT5b transporter throughout all light intensities, indicating slow gating mechanisms of EAAT5b.

**Figure 5:**
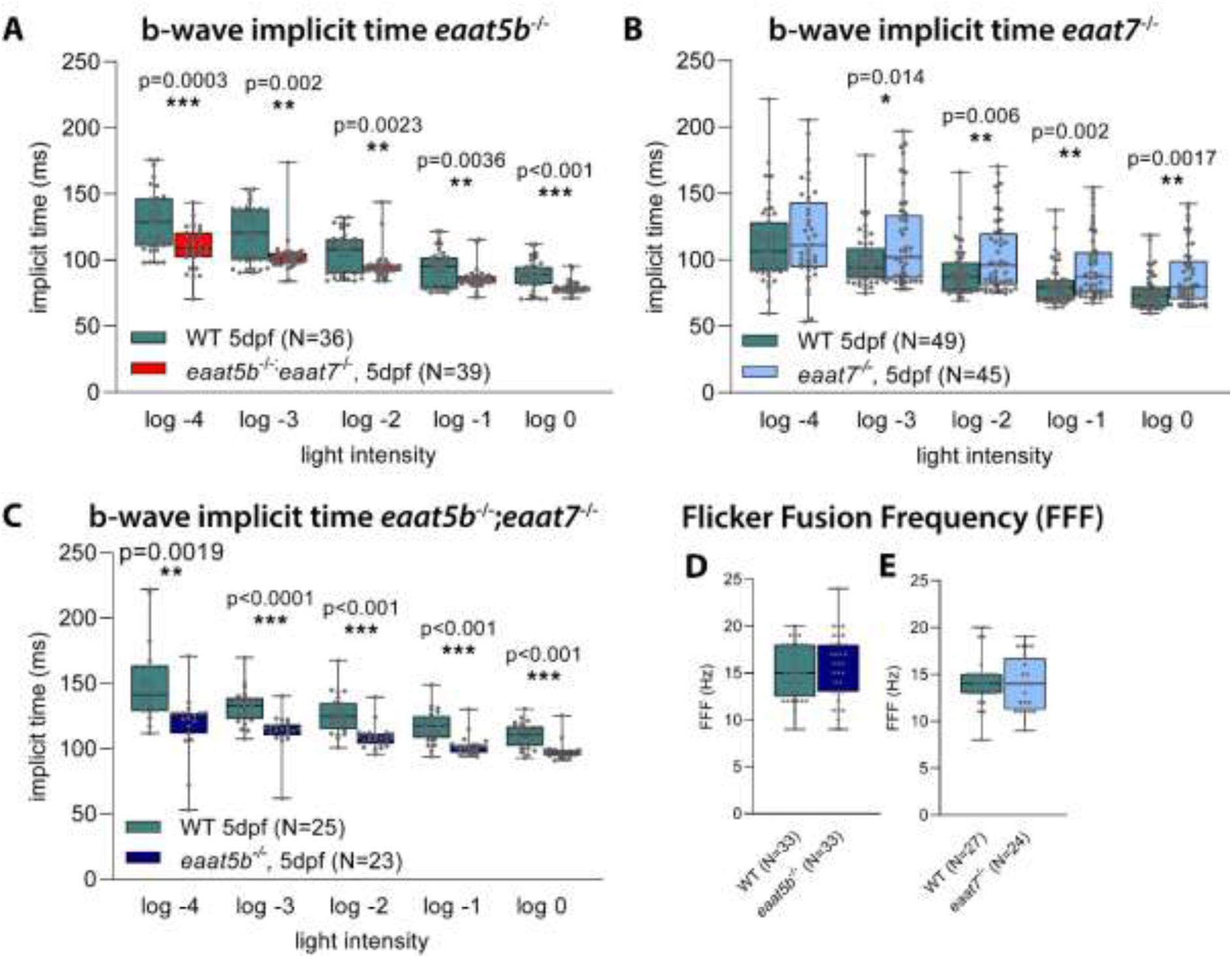
EAAT5b and EAAT7 differentially shape the ON-response but do not influence the flicker fusion frequency (FFF). The time from light onset to the initiation of the b-wave was quantified and plotted in box-and- whisker plots for *eaat5b* mutant (A), *eaat7* mutant (B) and double KO animals (C). Loss of EAAT5b significantly reduces the implicit time across all irradiances (A). Mutation in *eaat7* results in an increased implicit time (B). If both transporters are lost, the ON-response implicit time is decreased (C), alike in the *eaat5b* single mutant. Nevertheless, the slow gating mechanism of EAAT5b is not affecting temporal resolution of vision, as both, *eaat5b* (D) as well as *eaat7* mutant (E) animals display a FFF similar to WT.

Conversely, the ON-response initiation in *eaat7* mutants is delayed compared to WT animals for all light intensities except for very dim light stimuli (log −4) (Figure 5B), indicative for different gating kinetics between EAAT5b and EAAT7. The double KO *eaat5b*;*eaat7* resembles the single *eaat5b* mutant in showing a decrease in the ON-response implicit time for all light intensities (Figure 5C).

In order to further explore kinetic properties of the mutants, we assessed the flicker fusion frequency by ERG. This method measures the time for full recovery of the ERG response after a conditioning flash of light. Despite the differences in gating kinetics, no significant change in the flicker fusion frequency could be observed in neither *eaat5b*-/- nor in *eaat7*-/- (Figure 5D-E).

### Electrophysiological characterization of *dr*EAAT7 in HEK293T cells

We next expressed *dr*EAAT7 and *dr*EAAT5b fused to YFP transiently in HEK293T cells and tested surface insertion of *dr*EAAT7-YFP with confocal microscopy. Confocal images and fluorescence intensity plots of cell passing transects showed almost exclusive insertion of WT *dr*EAAT7-YFP into surface membrane, or in domains in close proximity (Figure 6 A). EAAT5b was only expressed at low levels, and we could not detect clear membrane staining (Supplementary Figure 4). Electrophysiological characterization of *dr*EAAT5b was not possible in mammalian cells. EAATs are secondary-active glutamate transporter and also anion channels (Fairman et al. 1995b; Machtens et al. 2015; Wadiche et al. 1995; Winter et al. 2012), and we therefore had to experimentally separate the two transport functions of *dr*EAAT7. Glutamate is co-transported into cells together with 3 Na^+^ and 1 H^+^, followed by the K^+^- bound re-translocation of the transporter back to the outward-facing state, and anion channel openings occur from intermediate conformations (Figure 6b) We tested *dr*EAAT7 mediated glutamate transport by whole cell patch clamping under conditions abolishing EAAT anion currents: cells were internally dialyzed with K^+^ gluconate and externally perfused with Na^+^ gluconate (Kovermann, Hessel et al., 2017, Watzke, Bamberg et al., 2001). Cells were clamped at a holding potential of −120 mV, and glutamate uptake was evoked by adding 5 mM to the external solution. Glutamate caused a reversible inward current in cells expressing zebrafish EAAT7 (Figure 6 C), but not in untransfected control cells (data not shown). Figures 6 D, E compare uptake currents mediated by zebrafish EAAT7 to human EAAT1 and rat EAAT4 uptake currents (n = 6/5/4). *h*EAAT1 represents an efficient glial glutamate transporter (Banner et al. 2002; Kovermann et al. 2017; Storck et al. 1992), and *r*EAAT4 is a prototypical glutamate transporter with low transport capability and high anion conductance (Kovermann, Machtens et al., 2010, Machtens, Kovermann et al., 2011, Melzer, Torres-Salazar et al., 2005). The uptake capability of zebrafish EAAT7 is similar to *r*EAAT4, (*p* = 0.332) but much smaller than *h*EAAT1 (*p* < 0.001, Figure 6 D, E).

**Figure 6:**
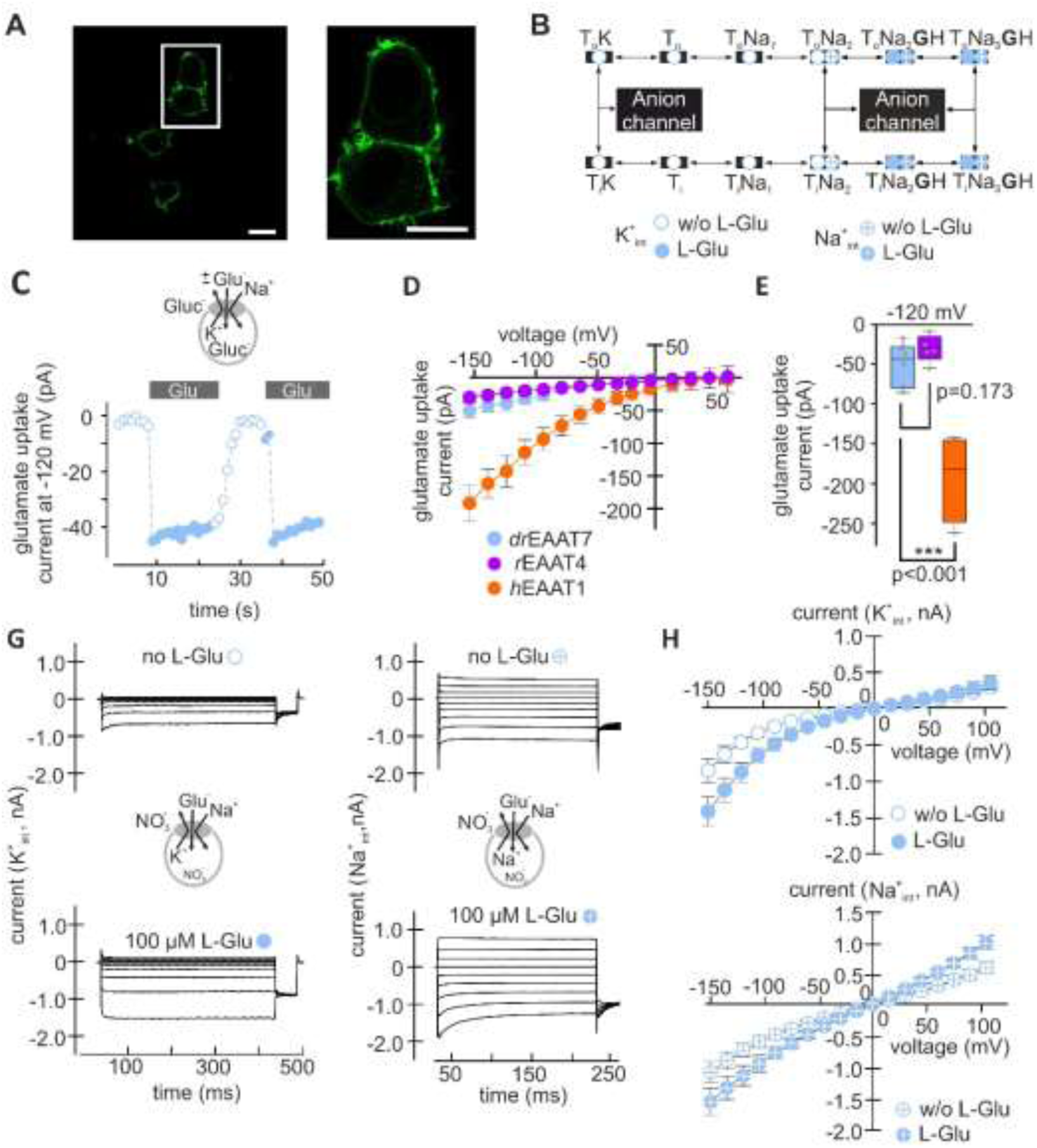
Zebrafish EAAT7 exhibits low transport capability and a dominating anion conductance. (A) Confocal microscopy indicates an almost exclusive insertion of the EAAT7-YFP fusion protein into the cell membrane (scalebars: 10 µm). (B) Scheme of the full transport cycle (empty circles) and of the Na^+^-exchange half-cycle (crossed circles) depict the transitions from transport mode to anion channel openings during the intermediate conformations between outward (o) and inward (i) facing conformations. (C) Whole cell glutamate transport currents in the absence of permeable anions at a holding potential of −120 mV over time. The applications of 5mM L-glutamate is indicated by grey bars (controls: open circles, with glutamate: closed circles). (D) Current voltage relationships of transport currents for *dr*EAAT7, *r*EAAT4, and *h*EAAT1, under ionic conditions as used in 6C (n = 6/5/4). (E) Boxplot of transport current amplitudes at −120 mV for *dr*EAAT7, *r*EAAT4 and the high-capacity transporter *h*EAAT1. (G) Representative whole cell currents in the presence of the permeable anion NO_3_^-^ under control conditions (top) and in the presence of 0.1 mM L-glutamate in the bath. Currents are shown for ionic conditions permitting the full transport cycle (K^+^_int_, left), and restricting the transporter conformations to the exchange half-cycle (Na^+^_int_, right). (H) Current voltage relationships for the whole cell anion currents shown in G for both conditions (n = 7/7).

The EAATs transport either Na^+^ together with glutamate and H^+^ or K^+^ in two half-cycles, often denoted as Na^+^- and K^+^ hemicycles (Figure 6 B). To record anion currents associated with both hemicycles (Machtens et al., 2015) we dialyzed cells with NO ^-^-based solutions supplemented with K^+^. Anion channel activity associated with the Na^+^-hemicycle were measured with Na^+^ as main internal cation (Figure 6 G, H). Application of glutamate increased currents at negative holding potentials under uptake conditions (Figure 7 H, −150 mV, *P* = 0.016 (K^+^*_int_*, n = 7), but not under exchange conditions *p* = 0.107 (Na^+^*_int_*, n = 7). In the presence of K^+^*_int_*, this increase was restricted exclusively to negative holding potentials (+75 mV, *p* = 0.515), while with Na^+^*_int_* significant current increases were observed only at positive holding potentials (+75 mV, *p* = 0.017). We conclude, that *dr*EAAT7 is a functional glutamate transporter with a dominating anion conductance.

## Discussion

In darkness, photoreceptors tonically release the excitatory neurotransmitter glutamate. Upon light stimulation, photoreceptors hyperpolarize leading to a reduction or cessation of glutamate release. ON-bipolar cells react to the decreasing glutamate concentration in the synaptic cleft by depolarization. This paradoxical inhibition by the quintessential excitatory neurotransmitter glutamate is thought to be mediated by the G-protein coupled mGluR6-TRPM1 pathway and conserved across vertebrates (Morgans et al. 2009; Audo et al. 2009; Huang et al. 2012). A number of studies conducted on different vertebrates suggest an additional non-metabotropic glutamatergic mechanism to be involved in ON-bipolar cell activation, mainly in teleosts (Nawy and Copenhagen 1987; Grant and Dowling 1995) but also in mammals (Tse et al. 2014). The described glutamate-gated chloride current I_Glu_ hyperpolarizes ON-bipolar cells in darkness, in the presence of glutamate. Decreasing glutamate concentrations in the synaptic cleft upon light reduced this current and caused by, depolarize ON-bipolar cells. I_Glu_ was shown to be sensitive to the non-specific EAAT blocker TBOA, implying to be generated by a glutamate transporter associated with high Cl^-^ conductance (Grant and Dowling 1996).

It is still not clear, which EAAT family members are responsible for glutamate reuptake in the retina and the details of the interplay between EAATs with the well-defined mGluR6 pathway remained equally unresolved. Previous research proposes a role for EAATs in cone signaling, while mGluR6 mainly functions in rod signaling in the giant Danio (Wong et al. 2005a). Other studies suggest a spectral difference between mGluR6 and EAAT signaling. mGluR signaling is thought to mainly mediate short wavelength photopic signaling, while calculations of the spectral sensitivity of the non-metabotropic signaling revealed it to be distributed across the visible spectrum (Saszik et al. 2002; Nelson and Singla 2009).

In the present study, we identified EAAT5b and EAAT7 as the glutamate transporters that mediate direct synaptic transmission in the outer retina. EAAT5b and EAAT7 were found to be expressed on dendritic tips of ON-bipolar cells and therefore in the expected localization for mediating these hyperpolarizing Cl^-^ currents in presence of glutamate, described by (Nawy and Copenhagen 1987; Grant and Dowling 1995, 1996; Connaughton and Nelson 2000; Nelson and Singla 2009). EAAT5b and EAAT7 co-localize on ON-bipolar cell dendritic terminals that are in contact with all four cone subtypes but only a limited number of rod spherules, indicating an important function in cone signal transmission. Expression of EAAT5b and EAAT7 does not indicate a spectral difference in the involvement of glutamatergic signaling pathways activating ON-bipolar cells as localization does not discriminate between synapses of different cone types.

In order to assess the function of these two EAATs in synaptic transmission, we generated single knockout animals and double mutants. No apparent changes in retinal morphology and localization of various synaptic markers were apparent. Functional analysis of the two glutamate transporters in the retina were performed by ERG recordings, focusing on the b-wave as a read-out of ON-bipolar cell depolarization. It is important to note that at the developmental stages used (5 dpf), rods do not significantly contribute to the ERG, hence the measured responses reflect pure cone responses (Bilotta et al. 2001) (Venkatraman et al. 2020). Lack of EAAT5b causes a slight decrease in the b-wave amplitude for medium light intensities, indicating that EAAT5b contributes to hyperpolarization of ON-bipolar cells in darkness, albeit to a small extent. *eaat5b/eaat7* showed a robust reduction of the b-wave, proving a redundant involvement of these two glutamate transporter in the ON-response of the cone pathway. The remaining smaller b-wave in the double mutant argues for a joined contribution of non-metabotropic (EAAT) and metabotropic (mGluR6) signaling in the generation of the photopic ON-response. Interestingly, studies in the adult Giant Danio, a closely related species, presented evidence for a much larger contribution of EAATs on the cone ON-response (Wong et al. 2005a; Wong et al. 2005b).

*eaat5b* knockout animals show a significant decrease in the ON-response implicit time, meaning that initiation of ON-bipolar cell depolarization is accelerated in mutants. This observation precludes the possibility of EAAT5b being a high capacity transporter, as a lack of such a high capacity transporter would cause a delay of the ON-response caused by remaining cleft glutamate even upon a light stimulus. The analysis of the ERG components by (Nelson and Singla 2009) revealed that the non-metabotropic ON-response consists of a short negative deflecting wave followed by the bigger positive deflecting b-wave. This post-photoreceptor a-wave is likely caused by the Cl^-^ current that is reduced when glutamate levels decrease at light on (Nelson and Singla 2009). The mammalian retina-specific EAAT5 has been associated with slow gating mechanisms (Gameiro et al. 2011), and it is therefore tempting to speculate that zebrafish EAAT5b also exhibits slow activation and deactivation and that the post-photoreceptor a-wave is generated by such slowly activating and deactivating EAAT5b Cl^-^ currents. Loss of EAAT5b would cause the abolishment of this post-photoreceptor a-wave allowing ON-bipolar cells to depolarize faster, as seen in *eaat5b* mutant animals. Remarkably, EAAT5 is presynaptically expressed in the mouse retina and is the source of autoinhibition of rod bipolar cells (Lukasiewcz et al. 2021)(Gehlen, Aretzweiler et al., 2021).

EAAT7, lacking in mammalian retinas, is be associated with fast kinetics. *eaat5b*-/-;*eaat7*-/- animals displayed the same acceleration in the ON-bipolar cell depolarization as *eaat5b* single mutants, supporting the hypothesis of the slow initiation of the ON-response being caused by the short hyperpolarization of ON-bipolar cells at light on (post-photoreceptor a-wave). The observed kinetics of the ON-response are especially remarkable considering that ON-bipolar cell activation in double mutants is solely mediated by metabotropic signaling, which is generally assumed to be slow due to the involvement of second messengers (G protein signaling). Nevertheless, our kinetic analysis demonstrates that mGluR6 mediated depolarization of ON-bipolar cells is of a faster time frame than EAAT5b mediated ON-bipolar cell depolarization. The slow kinetics of EAAT5b has no influence on temporal resolution of vision, as the flicker fusion frequency (FFF) in *eaat5b* and *eaat7* KO animals are comparable to the one in WT animals, indicating that other mechanisms are the rate limiting factors of FFF. Interestingly, the single EAAT5 protein was found to improve the temporal resolution of the mouse retina, via a presynaptic mechanism in bipolar cells (Gehlen et al. 2021).

Taken together, EAAT5b and EAAT7 are the two transporters that mediate the previously described glutamate gated Cl^-^ current in zebrafish ON-bipolar cells. Thus, in darkness when glutamate is maximally released, the Cl^-^ flux through EAATs renders ON-bipolar cells hyperpolarized. At light on, termination of glutamate release by photoreceptors causes ON-bipolar cells to depolarize due to the termination of the EAAT mediated Cl^-^ current and the opening of the cation conducting TRPM1 channel mediated via mGluR6 signaling. We showed that EAAT5b and EAAT7 are postsynaptically expressed in ON-bipolar cell synapses contacted by all cone subtypes. Loss of these EAATs leads to a diminished ON-response, proving their partially redundant involvement in direct synaptic transmission. Having a dual glutamatergic input system harbors several advantages. Rods seem mainly rely on metabotropic signaling which allows signal amplification. Furthermore, EAATs and mGluRs are thought to have different glutamate dose response functions. The demonstrated kinetic effect on the ON-response allows for an additional level of temporal regulations. Salamander EAAT5 expressed in *Xenopus* oocytes was shown to have a glutamate dose response over 4 log units (Eliasof et al. 1998), while mGluR6 glutamate dose response saturates at 2 log units (Shiells and Falk 1994). And last but not least, mGluRs are not only involved in signal transmission but are key molecules in mediating proper localization of synaptic proteins (Tummala et al. 2014; Tummala et al. 2016).

## Acknowledgements

All authors would like to thank Martin Walther and Kara Kristiansen for excellent technical assistance and animal care taking. Further acknowledgements go to Andrea Gmür for support with Flicker ERG analysis and to Dr. Igor Delvendahl for his help with ERG kinetics analysis. We are also grateful to Drs. Christian Mosimann and Martin Jinek for kindly providing us with Cas9 protein and Dr. Shoji Kawamura for sharing his transgenic lines.

This work was funded by the Swiss National Science Foundation (310030_204648 and 310030_200376)

## Supplemental Information

**Supplemental Figure 1:**
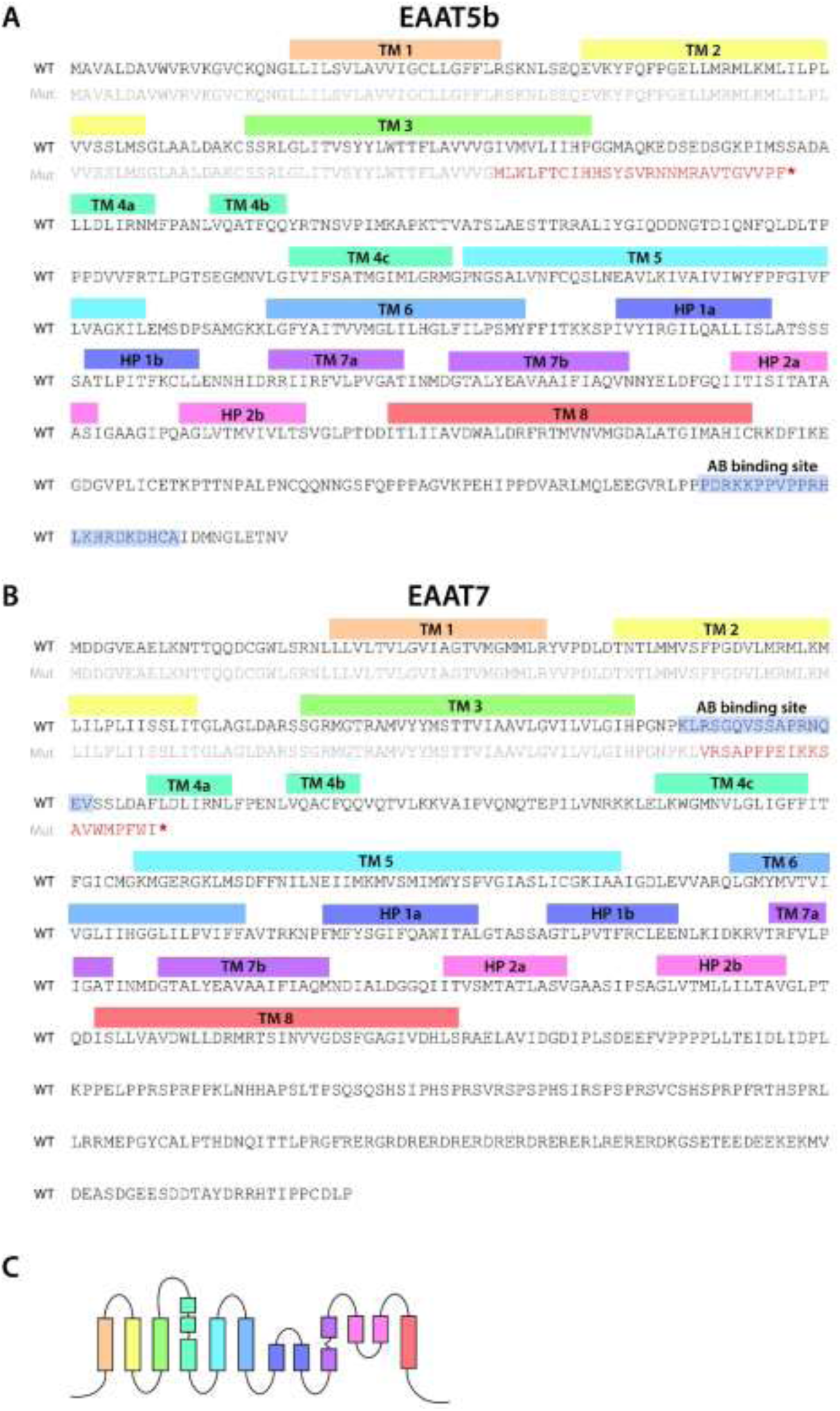
WT and mutant protein sequences with labeled domains. WT protein sequences for EAAT5b (A) and EAAT7 (B) are shown in black font and mutant sequence in gray. +79 bp indel in *eaat5b* results in a premature stop codon (red asterisk) between TM 3 and 4. Antibody binding site is at the C-terminus of the protein (A). 4 bp deletion in *eaat7* results in a premature stop within TM 4. Antibody binding site lies between TM3 and TM4 and is disrupted by the mutation (B). WT EAAT protein structure (C). TM: transmembrane domain; HP: hair pin; AB: antibody.

**Supplemental Figure 2:**
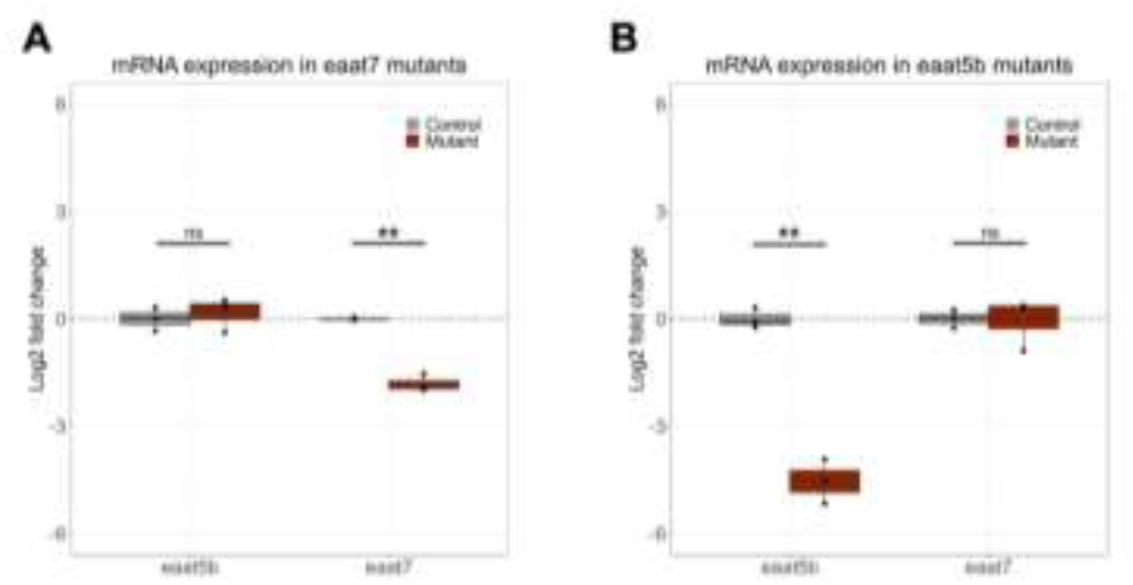
mRNA transcript levels of eaat5b and eaat7 in eyes of eaat7-/- (A), and eaat5b-/- (B) 5 dpf larvae relative to their wild type siblings. Transcripts were measured by RT-qPCR and normalized to rpl13a and actb2. Data are represented in box-and-whisker plots with an interquartile range from first to third quartile and the median represented by the line within the boxes. Black dots represent individual sample means. Significance levels: ***p < 0.001, **p < 0.01, *p < 0.05, ns = not significant (p > 0.05), Welch two sample unpaired t-test.

**Supplemental Figure 3:**
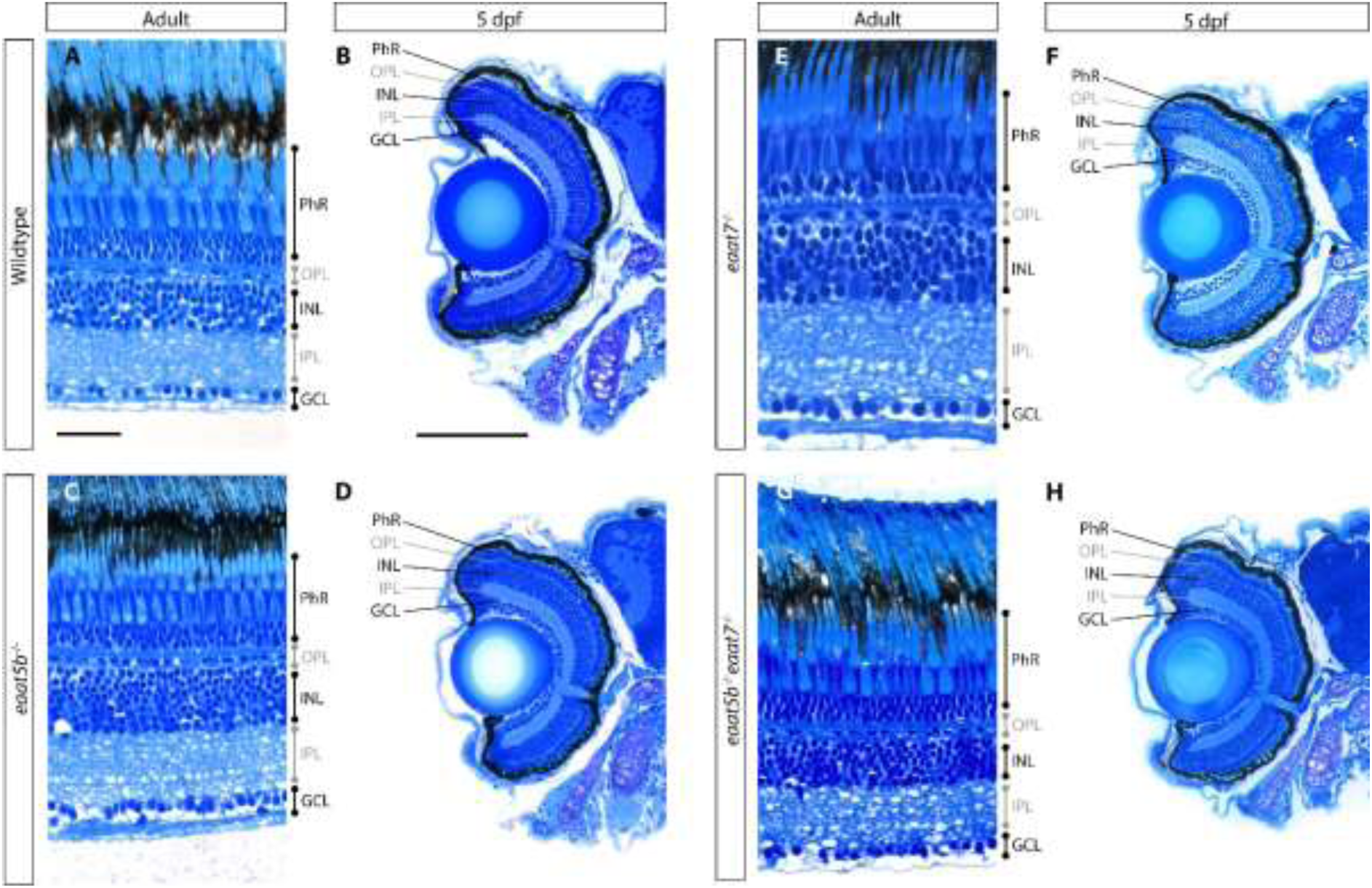
Normal retinal morphology in *eaat5b*-/-, *eaat7*-/- and *eaat5b*-/-;*eaat7*-/- animals. Histological sections of adult (A, C, E, G) larval (5 dpf) retinas (B, D, G, H) reveal that the morphology of the retina seems unaffected in *eaat5b* (C, D), *eaat7* (E, F) and *eaat5b*;*eaat7* double KO (G, H) animals. Scale bar in A is 30 µm and also corresponds to C, E and G, scale bar in B is 100 µm and also corresponds to D, F and H. PhR: photoreceptor; OPL: outer plexiform layer; INL: inner nuclear layer, IPL: inner plexiform layer; GCL: ganglion cell layer.

**Supplemental Figure 4:**
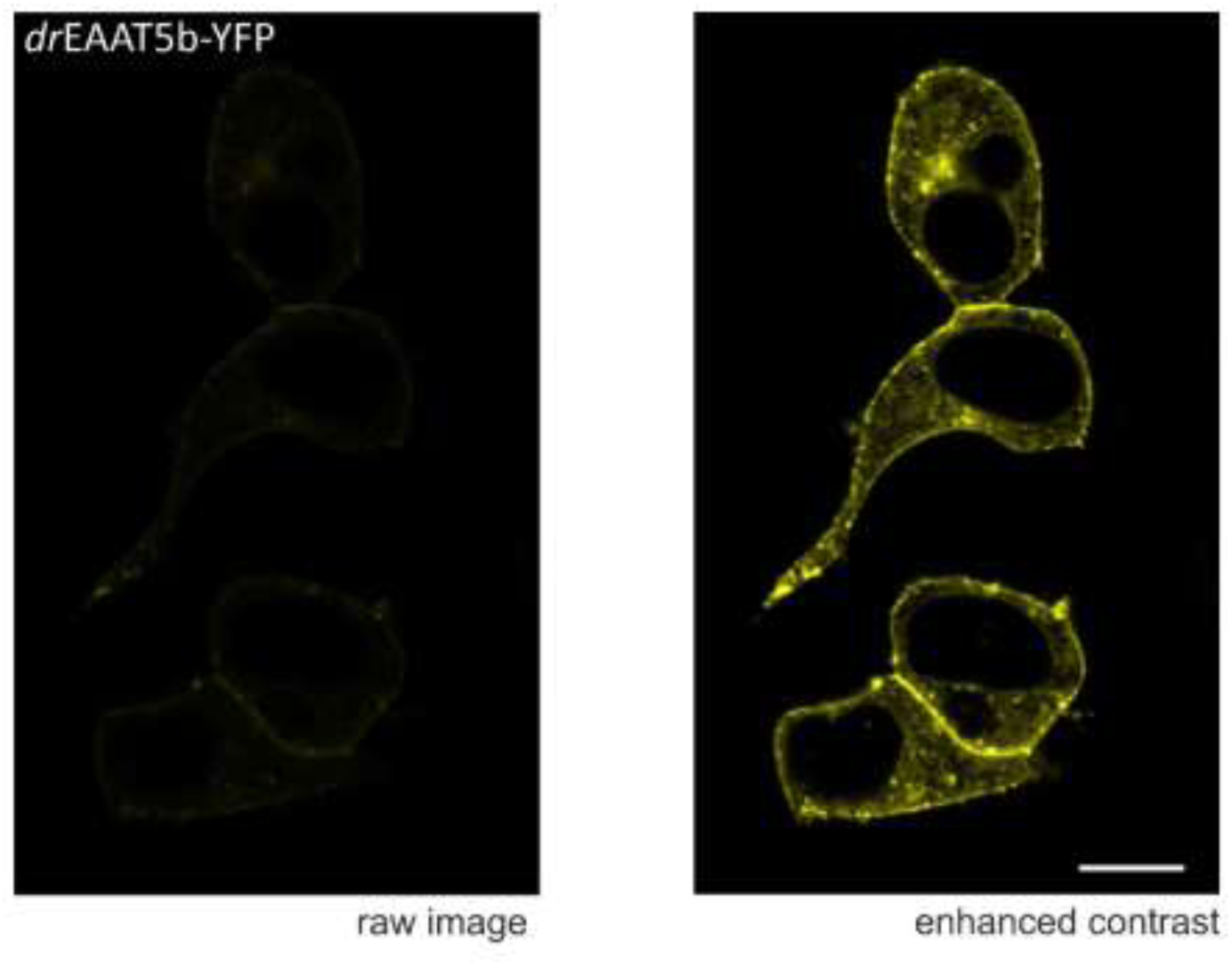
Zebrafish EAAT5b is weakly expressed in HEK293T and does not insert into the cell membrane. The figures show confocal microscopy recordings from HEK293T from cells expressing *dr*EAAT5b- YFP fusion protein. The left figure shows the original raw image, under the same recording conditions as for *dr*EAAT7 (Figure 6), and right figure shows increased contrast. EAAT5b is accumulated in the cytosol, and in patchy regions next to the cell surface (scale bar: 10 µm). Attempts to characterize EAAT5b with the patch clamp technique failed. No whole cell currents were detected (data not shown).

## Notes

### Competing Interest Statement

The authors have declared no competing interest.

